# From Bayes to Darwin: evolutionary search as an exaptation from sampling-based Bayesian inference

**DOI:** 10.1101/2023.05.29.542733

**Authors:** Márton Csillag, Hamza Giaffar, Eörs Szathmáry, Mauro Santos, Dániel Czégel

## Abstract

Building on the algorithmic equivalence between finite population replicator dynamics and particle filtering based approximation of Bayesian inference, we design a computational model to demonstrate the emergence of Darwinian evolution over representational units when collectives of units are selected to infer statistics of high-dimensional combinatorial environments. The non-Darwinian starting point is two units undergoing a few cycles of noisy, selection-dependent information transmission, corresponding to a serial (one comparison per cycle), non-cumulative process without heredity. Selection for accurate Bayesian inference at the collective level induces an adaptive path to the emergence of Darwinian evolution within the collectives, capable of maintaining and iteratively improving upon complex combinatorial information. When collectives are themselves Darwinian, this mechanism amounts to a top-down (filial) transition in individuality. We suggest that such a selection mechanism can explain the hypothesized emergence of fast timescale Darwinian dynamics over a population of neural representations within animal and human brains, endowing them with combinatorial planning capabilities. Further possible physical implementations include prebiotic collectives of non-replicating molecules and reinforcement learning agents with parallel policy search.

## Introduction

Learning and memory are a major focus in the behavioural sciences (e.g. anthropology, biology, cognitive science, economics, psychology, political science, sociology) and in computer science. At its most basic level, learning is the ability of an entity to acquire knowledge through experience or imitation, via non-associative or associative learning. Non-associative learning takes the form of habituation (learning not to respond to repeated or irrelevant stimuli), and sensitization (learning to respond to relevant stimuli). Associative learning is the establishment of an association between a stimulus and a response. Learning can also be a matter of knowledge representation, i.e. a surrogate for the state of the word, and the handling of uncertainty. “Memory in biological *and artificial* systems always entails learning (the acquisition of information) and … learning implies retention (memory) of such information”(1) (p. 751; our addition in italics).

General ideas from probability theory and statistical inference from a Bayesian perspective, which views learning as the updating of probabilistic beliefs, have provided a bridge between learning theory and practice. Bayesian inference is widely used in machine learning (2). In recent years, several scholars have pointed out the remarkable analogy between Bayesian inference and Darwinian replicator dynamics in asexual populations (3–7). From an evolutionary perspective, these dynamical-structural analogies suggest that whole populations of replicators can perform Bayesian computations even though they are not themselves units of selection, i.e. without community-level selection (7). Simple systems of Darwinian replicators can implement basic statistical inference algorithms, such as Bayesian inference (3–5), filtering in hidden Markov models (8), and inference in hierarchical models (9). In addition, evolution exhibits examples of emergent memory across scales, formalized as autoassociative networks (10). Such memory storage and adaptive retrieval aid development when implemented by gene regulatory networks (11–13). At a much larger temporal and spatial scales, this capacity also provides adaptive explanations at the ecosystems level when implemented by evolutionary ecological dynamics among interacting species (14).

Environments are noisy and change over various timescales - extracting relevant information from such environments allows prediction of future environments and is essential for organismal survival and reproduction, as well as adaptive behaviour by artificial agents (15–17). However, from a genetic perspective, learning is a form of phenotypic plasticity, and in some cases, this plasticity can be maladaptive, i.e. when organisms face increasingly stochastic change (18). We note here that the first publication linking learning (plasticity) and evolution was by the American psychologist James Mark Baldwin in 1896 (19). He was interested in the idea that organisms can learn to adapt to a new environment they encounter, and that those who learn quickly have an evolutionary advantage. Over many generations, these learning behaviours could become innate: “When learning guides evolution” was the title of a commentary in *Nature* by Maynard Smith (20). Our emphasis here is radically different. What we propose is a mechanism for the *emergence* of Darwinian evolution via Bayesian computations, as a possible consequence of the algorithmic equivalence between replicator-mutator dynamics and filtering in hidden Markov models (see Figure 1B and C, as well as ref. (7) for details).

**Figure 1.**
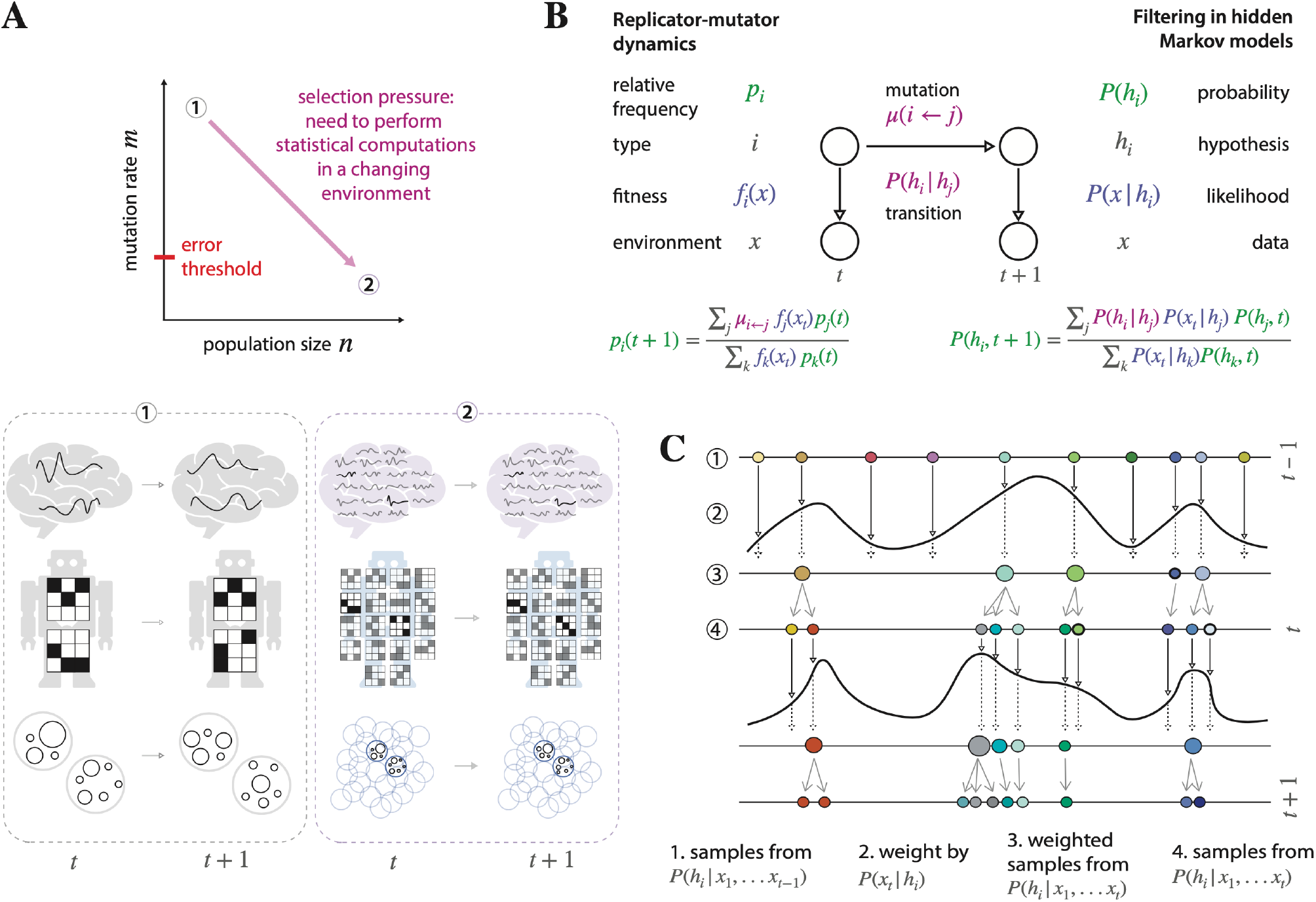
A Bayesian objective provides a possible route towards the emergence of self-replication in brains, artificial agents, and prebiotic molecular collectives. A) When collectives of representational units (neural ensembles, policy representations, or prebiotic molecular systems) are selected for estimating the Bayesian posterior of changing environments, inheritance of information between units/particles within collectives can emerge. Continued selection for Bayesian inference at the level of collectives leads to Darwinian evolutionary dynamics of a large population of high-fidelity replicators within collectives, a system surpassing Eigen’s error threshold. Even though never selected for it, such replicator populations can perform efficient combinatorial search when they unfold over longer timescales. B) Computational level equivalence between replicator-mutator dynamics and the dynamics of the filtering distribution in Hidden Markov Models. C) Algorithmic equivalence between a model of population genetics and particle filters, an efficient sampling-based approximation of the filtering distribution in Hidden Markov Models. Together, B and C entail a partial unification of Bayesian and Darwinian dynamics, which forms the basis of the scenario depicted in A.

The partial unification of replicator dynamics and Bayesian computation makes it possible to bridge the two domains (6,7). In practice, it opens up the possibility of hybrid architectures and mechanisms that exhibit both Darwinian and Bayesian properties (21,22). We suggest that when collectives are under selection for their ability to perform accurate statistical inference over changing environments, they can undergo a Darwinian transition, similar to a critical step during the origin of life, in which highly accurate self-replication of information emerges, providing unlimited hereditary potential (23,24). Such a transition involves an adaptive trajectory that begins with a few cycles of noisy information transfer between two units. This leads to true Darwinian evolution with high-fidelity information replication among a population of units over many generations. As a result, the population gains the ability to maintain and cumulatively improve complex adaptations, formalised as improved performance on combinatorial optimization tasks.

In biological evolution, for example, new functions often appear through exaptation (preadaptation in the older terminology), as discussed in detail in ref. (25). In this work, we show that the function to perform combinatorial optimization can be exapted from a faculty of Bayesian inference. In more technical terms, this means that the evolved parameters that allow efficient Bayesian inference also happen to allow combinatorial optimization—in a rudimentary form. A didactic analogy is the role of evolution of feathers in flight. Feathers first appeared in dinosaurs to keep them warm (26). They were later found to be useful for simple gliding and eventually for real flight. This was made possible by direct selection for the new function from the humble original state that served the old function. In the same way, the mastery of combinatorial optimization requires further evolution of the parameters.

In animal brains, relevant representational units may be neural ensembles that scale from a few loosely functionally connected ensembles to a large number that can support high-precision copying of information between them, as suggested by the hypothesis of Darwinian neurodynamics (27–31). From a cognitive perspective, Darwinian neurodynamics is based on the idea that evolutionary search in the space of candidate solutions to complex (combinatorial, non-differentiable) problems might be possible in the brain in real time, outperforming simpler search mechanisms such as solitary and parallel stochastic search. Parallel search with information transfer between units is arguably the most effective strategy when the search space has a complex associated landscape, as discussed in ref. (28). One example is that of different activity patterns occupying topographically different locations in the corresponding brain area as candidate solutions (29,31). Note that Darwinian neurodynamics differs from the Neural Darwinism of Edelman (32) in that the former rests on the classical multiplication-heredity-variability conditions of evolutionary units (33), whereas the latter does not implement an evolutionary algorithm, but is purely selectionist and based on the idea that synapses or groups of synapses are differentially stabilised by reward.

In artificial agents, such as reinforcement learning architectures, a small number of appropriate architectural and hyperparameter choices could lead to information replication and Darwinian evolution over representations (34–37). We suggest that these choices include a modular representation of the environment, with interconnected and architecturally similar modules required for high-capacity information transfer and shared representational capacity. In all cases, redundancy can be considered a “smoking gun”: redundant cortical representations of action plans (38), or population-based *in silico* architecture design (36,39) must work against the selective forces of energetic and storage capacity costs. In a biological context, Darwinian neurodynamics could underlie learning by operant conditioning of combinatorially complex behavioral patterns.

The equivalence between Bayesian computation and Darwinian replicator dynamics extends to both Marr’s computational and algorithmic levels (40). At the computational level, the dynamics of relative frequencies in the discrete-time replicator-mutator equation is equivalent to the dynamics of the filtering distribution in hidden Markov models (HMM) (7,8). At the algorithmic level, the Darwinian mechanism of multiplication, inheritance and selection operating on a population is mapped onto a sampling-based approximation of the HMM’s filter distribution, called particle filters (6,7,41,42). A fundamental difference between the two domains is the parameter regimes that lead to “good” computations. In HMMs, the task is to infer the posterior (filtering) distribution accurately. Conversely, evolutionary dynamics is often considered and applied as an optimizer in large combinatorial spaces with many local optima. Here, we combine these two ideas and show that a continuous adaptive trajectory, from very low information transmission between two units to Darwinian evolution over many units, is achievable if selection comes from statistical inference of stochastic environmental dynamics.

## Model

We formulate a model that contrasts statistical inference with optimization within a single framework. Importantly, our population-based model spans the space from no information transfer between units/individuals to near perfect copying, and it sets the number of units participating in the collective computation as a free and potentially adaptive hyperparameter. The model is intended to be as agnostic as possible with respect to both specific representational choices (e.g., how and where information is stored and transmitted) and statistical inference or optimization tasks.

### NK landscapes

We choose NK landscapes (43–45) to represent both statistical inference and optimization tasks with tunable complexity. Although abstracted from evolutionary theory, we suggest that optimizing or sampling NK landscapes can provide a simple task-agnostic representation of combinatorial cognitive tasks with many non-equivalent locally good solutions. NK landscapes form a statistical class of combinatorial optimization problems over the space of binary strings of length *N*. Inspired by the genotype-phenotype-fitness maps in biological evolution, where multiple genes collectively determine phenotypic traits and thus fitness, NK landscapes assign a scalar fitness value Φto binary strings (genotypes) *g* = (*g*_1_,*g*_2_,…,*g*_*N*_)as a sum of *M* fitness components (phenotypic traits) *f*_*i*_, 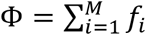. Each fitness component *f*_*i*_ is an independent random variable drawn from the uniform distribution over [0,1]. The complexity (ruggedness) of the landscape Φ(*g*)is parameterized by *K*, setting the *number of arguments* of each fitness component. *K* = 1 encodes a linear landscape, 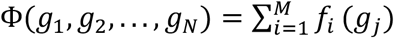 with a single global maximum and no other local maxima, whereas *K* > 1 describes rugged, nonlinear surfaces with multiple non-equivalent local maxima. This ruggedness is due to *epistasis*, i.e., the fact that each independent fitness component *f*_*i*_ is determined by multiple (*K* > 1) genes; the net effect of a single gene on fitness depends on other genes. (Note that this is a departure from the original notation by Kauffman (43), where *K* corresponds to *K* + 1 in our model, with no epistasis parametrized by *K* = 0.) Epistasis, defined here by the number of arguments of fitness components *f*_*i*_, entails a many-to-one mapping between genes and phenotypic traits. Pleiotropy, on the other hand, is the one-to-many nature of this map. For simplicity, we choose *NK* landscapes to have uniform pleiotropy in a statistical sense, each gene influencing exactly *K* fitness components by imposing *f*_*i*_ to depend only on *g*_*i*_ when *K* = 1 (no epistasis, no pleiotropy), and on *g*_*i*-1_,*g*_*i*_,*g*_*i*+1_when *K* = 3. Fitness component values *f*_*i*_ are sampled from the uniform distribution over the unit interval [0,1] independently for each of the 2^*K*^ combination of arguments. Consequently, a specific realization of an NK-landscape is defined by 2^*K*^*M* parameters. In our simulations, we set *N* = *M* = 20. Figure 2B was constructed by sweeping through all 2^*N*^ states. The dashed lines, corresponding to randomly reshuffled fitness assignments, are analytical approximations based on the definition of information gain, calculated as the number of better neighbors = *N* 2^−1^.

**Figure 2.**
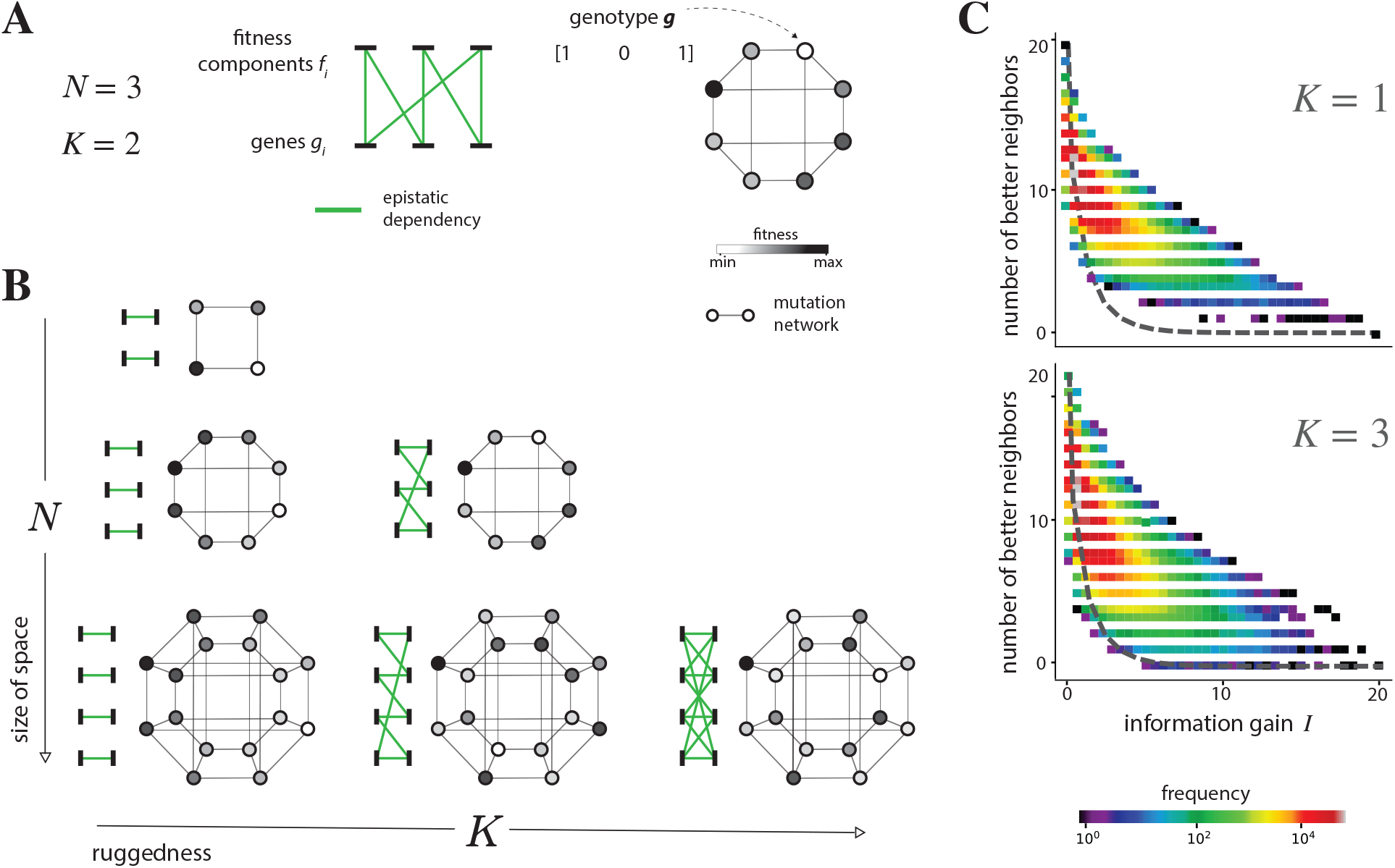
Tuneably rugged combinatorial surfaces model complex statistical inference and combinatorial optimization tasks. A) An example *N* = 3,*K* = 2 landscape containing 2^*N*^ genotypes, displayed as a mutation network in which nodes represent genotypes and edges represent single mutations between genotypes. Epistatic interactions between pairs (*K* = 2) of genes are shown in green: fitness components *f*_*i*_ depend on neighboring genes *g*_*i*_ and *g*_*i*+1_, with periodic boundary conditions. B) NK landscapes define statistical classes of scalar functions over binary strings, parametrized by the length of the string *N* and the ruggedness of the landscape *K*. C) The local fitness statistics of the landscape as a function of ruggedness *K*. The *x* axis represents the information gain value of a focal genotype, with the *y* axis representing the number of better (higher fitness) mutational neighbors of that focal genotype. Frequency of such *(x,y)* pairs over the landscape is indicated by color. The number of better mutational neighbors (local adaptive paths) is strongly determined by fitness when the landscape is linear (*K* = 1), but is highly variable when the landscape is rugged (*K* = 3). Yet local statistics of both linear (*K* = 1) and rugged (*K* = 3) landscapes are very different from landscapes with randomly reshuffled fitness assignments, with mean number of better neighbors shown by the dashed grey line.

In order to be able to compare NK landscapes across different *K* ruggedness parameters, we transform numerical fitness values to ordinal ones by ranking them in decreasing order, Φ(*g*)↦ *r*(*g*), and then we transform rank to a logarithmic scale, which we call *information gain I*(*g*)= ™log_2_(*r*(*g*)/*r*_*max*_) ∈ [0,*N*]. Information gain *I*(*g*)quantifies the amount of information (binary decisions or halving the remainder of the space) needed to find a genotype *g*′ that is at least as good as *g*.

Figure 2B and C illustrates the effect of the ruggedness parameter *K* on the local neighborhood of states: the number of mutational neighbors of *g*, that differ in exactly one bit from *g*, is plotted against information gain *I*(*g*). As the right panel of Figure 2B shows, rugged landscapes (*K* = 3) are still far from being random: fitness of neighbors are highly correlated, reflected by the fact that states with high fitness (*I*(*g*)≳ 2)have many neighbors with even higher fitness, compared to landscapes where fitness assignments are randomly reshuffled (dashed grey line). However, ruggedness has a strong effect on local neighborhoods: increasing *K* decreases the mean number of better neighbors but increases the variability of it across states. This is particularly apparent at the region of high fitness. In other words, adaptive mutational paths *from* high-fitness states become less available as ruggedness increases, and they also become strongly dependent on the state itself and not only its fitness. In the rest of the paper, we use NK landscapes with *K* = 1 and *K* = 3, representing linear and rugged surfaces, respectively.

### Statistical inference and optimization tasks

We define the classes of tasks that we will refer to as *statistical inference* and *optimization* below. In both cases, we exponentialize fitness values to make them positive, which allows the exponentialized fitness to be interpreted as the expected number of offspring, while preserving the complexity of the problem by using a monotonous transformation. Note that information gain *I*(*g*)and the local neighborhood statistics shown by Figure 2B are invariant under this (and any monotonous) transformation.

The *statistical inference* task is defined here as follows. Starting from a prior distribution *P*_/prior_(*g*)∝ exp(αΦ_/prior_(*g*)), the dynamics unfolds on the likelihood (of the environment *e*) *P*_likelihood_(*e*|g)∝ exp(*τ*αΦ_likelihood_(*g*))to compute the posterior distribution,

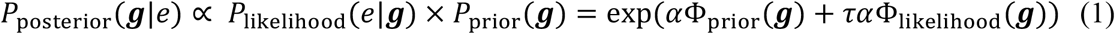

with Φ_/prior_ and Φ_likelihood_ being different realizations (different samples of the {*f*_*i*_} fitness component values) of NK landscapes with the same parameters *N* and *K. Optimization*, on the other hand, is simply climbing a single landscape, which we choose to be exp(*τ*αΦ_likelihood_(*g*))for maximal comparability. Intuitively, statistical inference entails an optimal integration of the past (prior) and the present (likelihood), which is well approximated by the population when evolution on the likelihood landscape is neither too fast nor too slow, and neither too directed nor too explorative. Furthermore, it is always computed at the transient regime due to the importance of remembering the initial condition encoded by the prior.

Another difference between the two tasks is how performance is evaluated. In the case of statistical inference, the task is to compute a probability distribution *P*_posterior_ over all 2^*N*^ genotypes. Accordingly, we quantify performance on the statistical inference task as the accuracy *A* of the population-based estimation of the true (exact) Bayesian posterior *P*_posterior_, measured as the Kolmogorov-Smirnov (KS) distance between the population-based estimate and the exact posterior. Optimization, on the other hand, consists of a search for the maximum information gain in the population, denoted by *I*_max<_. Note that both Bayesian inference accuracy *A* and optimisation performance *I*_max<_ are invariant under any monotonous transformation of fitness Φ, justifying their use as simple performance measures for both tasks.

Accuracy *A* is computed as follows. We compare two information gain distributions, the exact Bayesian posterior *P*_posterior_(*I*),constructed from all 2^*N*^ values, and the population-based estimate of it, *P*_population_(*I*),based on *n*_action_ values. Accuracy *A* is calculated as the Kolmogorov-Smirnov (KS) distance between these two empirical distributions, given by the maximal difference of their cumulative density function across *I*.

### Dynamics/algorithmic solution to the two tasks

The dynamics, similar to a Moran (46) model in population genetics and as particle filters or sequential Monte Carlo algorithms in Bayesian statistics, is the following. At every timestep, called generation, the population of *n* individuals, each represented by their genotype *g*, is resampled according to the individuals’ fitness/likelihood

*P*_likelihood_ = exp(*τ*αΦ_likelihood_(*g*)), and then mutated. Thus, a generation consists of *(i)* selecting an individual with genotype *g* for reproduction with probability proportional to its *P*_likelihood_, *(ii)* mutating it, *(iii)* adding it to the pool of the next generation, *(iv)* repeating *(i)-(iii) n* times. In our model, mutation is simply implemented as a point mutation (flipping each bit) with probability *m*, called the (per bit) mutation rate. Since individuals are represented by binary strings of length *N*, the per individual mutation rate is *Nm*; the expected total number of mutations in the population in a single generation is *n*N*m*.

To generate the population representing the *prior* distribution, exp(αΦ_/prior_(*g*)), we *independently* sample each particle via a Markov chain Monte Carlo (MCMC) algorithm. A single particle corresponds to the 1000th step (stationary regime) of a Metropolis-Hastings algorithm with proposal distributions set by point mutations.

Since the dynamics is assumed to be fully internal to the animal, agent, or collective, its parameters, mutation rate *m* and population size *n* are evolvable: on a timescale that is longer than that of the internal dynamics (e.g. genetic evolutionary timescales versus millisecond-timescale statistical computations in animal brains), the evolutionary parameters *m* and *n* follow an adaptive path on a cost surface given by their (average) performance in statistical inference or optimization. We do not model parameter evolution directly here, instead, we compare systems with different parameters (i.e., we evaluate the cost surface over the parameters) and assume that populations follow, at least approximately, an adaptive path on this cost surface.

Since the population size *n* serves as the sample size in estimating the Bayesian posterior, varying *n* introduces a bias in Bayesian accuracy *A*. We factor out this bias by introducing an *action selection* step when constructing population-based estimates of the posterior distribution. Action selection corresponds to uniformly selecting a *single* particle from the population of size *n* at the end of each run; then constructing the sample-based estimation of the posterior from *n*_action_ independent runs, each contributing one sample.

Another parameter that the animal, agent, or collective, can optimize is the way it breaks down the task to subtasks, each corresponding to one generation of the dynamics above. This is a strong deviation from conventional evolutionary models since it assumes that there exists a collective that can make an adaptive decision regarding this partitioning. On one end of the scale, it imposes a fitness landscape *P*_likelihood_(*e*|g)∝ exp(*τ*Φ_likelihood_(*g*))and performs the computation (either statistical inference or optimization) in a single timestep (generation). The other end of the scale is *T* >> 1 generations, which necessitates a proportionally less selective fitness landscape *at each generation*, exp(*τ*Φ_likelihood_(*g*))/*T*), so that the full computation, corresponding to an iterative update *T* times, provides an unbiased estimate of the posterior,

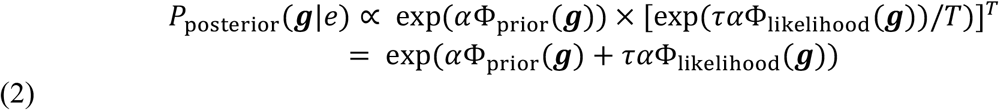

In our analyses below, we use the more intuitive parameter *τ*α/*T* = *s*, called selection strength, instead of the number of generations *T* directly. Correspondingly, the likelihood fitness landscape, parameterized by *s*, is given by exp(*s*Φ_likelihood_(*g*)). *τ*, on the other hand, is defined by the task itself, interpreted as the ratio of importance of the likelihood (present) and the prior (past). Figure 3A illustrates the relationship between parameters *s*,α,*τ*, and *T*; Table 1 presents an overview of all parameters, as well as their numerical value used in our simulations.

**Table 1.** lists the simulation parameters that are used to generate the results of Figure 4 and 5.

| symbol | parameter description | comment | evolvable? | Fig. 4. | Fig. 5. |
| --- | --- | --- | --- | --- | --- |
| $N$ | N parameter of NK landscape | number of binary genes that encode a genotype | | 20 | 20 |
| $K$ | K parameter of NK landscape | ruggedness | | 1, 3 | 3 |
| $\alpha$ | scaling parameter of the prior landscape | also called the *strength* of the prior | | 100 | 100 |
| $\tau$ | effective timescale of likelihood integration | strength of the likelihood $sT$ divided by the strength of the prior $\alpha$ | | 1, 10 | 1, 10 |
| $m$ | mutation rate | per bit mutation rate | evolvable | 0.001,...,0.5 | 0.001,...,0.5 |
| $s$ | selection strength | per generation selection strength; total strength of the likelihood is given by $s$ times the number of generations $T$; | evolvable; although the total strength of the likelihood $sT$ is fixed, $s$ is evolvable and it sets the number of generations $T$ | 0.5,...,50 | 0.5,...,50 |
| $n$ | population size (number of particles) | | evolvable | 100 | 2, 5, 10, 20, 50, 100 |
| $n_{\text{actions}}$ | number of samples used to estimate the posterior | at the end of each run, a *single* particle is sampled ("action selection"), the posterior is estimated based on $n_{\text{actions}}$ runs | | 100 | 100 |

**Figure 3.**
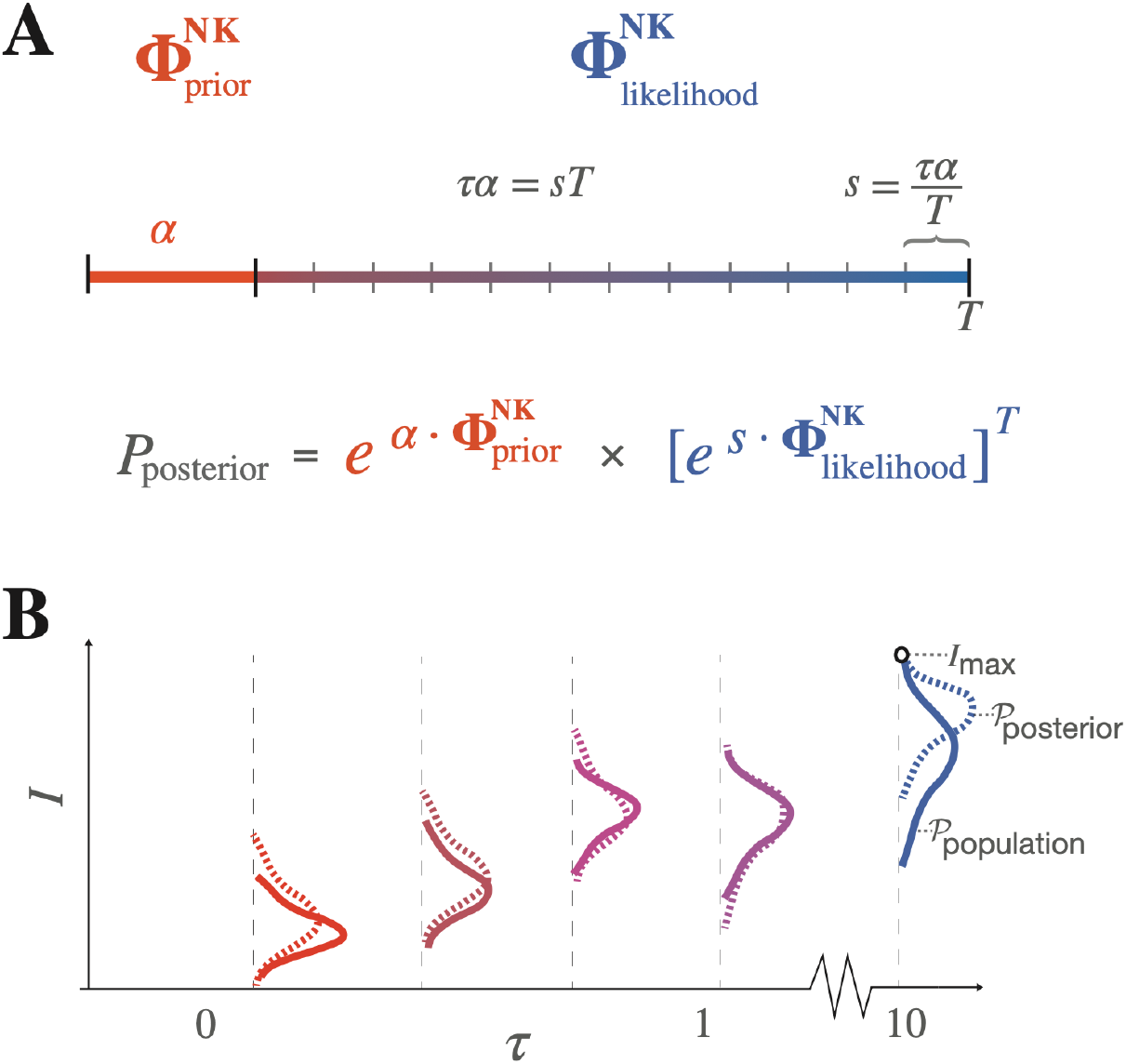
Bayesian inference and optimization in a single framework. A) Bayesian inference entails the integration of the prior and the likelihood, each represented by a particular realization of an NK landscape, 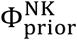 and 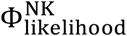. The strength of the prior and the posterior is set by α and *sT*, respectively, with *τ* = *sT*/α parametrizing the relative strength imposed by the likelihood versus the prior. The integration of the likelihood is adaptively subdivided into *T* computational steps, or generations, each experiencing a selection pressure *s*. B) We refer to the transient (*τ* = 1) regime as Bayesian inference, and the approximately stationary (*τ* = 10) regime as combinatorial optimization. The two tasks also differ in how performance is evaluated: optimization performance is measured by the maximum information gain *I*_max<_ of the population, while the accuracy *A* of Bayesian inference is evaluated by comparing the *distribution* of information gain values in the population *P*_population_ with the exact posterior distribution *P*_posterior_, measured by the Kolmogorov-Smirnov distance between them, *A* = KS(*P*_population_,*P*_posterior_).

For statistical inference tasks, we set *τ* = 1, which makes the importance of the prior, exp(αΦ_/prior_(*g*)),equal to that of the likelihood, exp(*τ*αΦ_likelihood_(*g*)),since Φ_/prior_ and Φ_likelihood_ are both NK landscapes with the same parameters *N* and *K*. For optimization tasks, on the other hand, we set *τ* = 10, approximately evaluating long-term dynamics on a single landscape exp(*τ*αΦ_likelihood_(*g*)). As before, we model both tasks and the corresponding dynamics in a unified framework wherever possible, which includes setting the initial population to be an unbiased sample of the prior, exp(αΦ_/prior_(*g*)). This is important for statistical inference as it evaluates the integration of the prior and the likelihood against an optimal one given by the true posterior, and it also guarantees that all statistics between the initial population and the likelihood landscape is fixed across tasks - i.e., there is no preadaptation for optimization.

Overall, we propose this as a minimal, proof of principle model that unifies Bayesian inference and Darwinian dynamics in a single framework to investigate adaptive hyperparameter trajectories over both Bayesian and optimization objectives. Most importantly, we *(i)* choose to be task agnostic with no explicit model of what the genotypes represent, using only a statistical model of the fitness or likelihood function and *(ii)* we evaluate performance in a way that is invariant over any monotonic transformation of fitness or likelihood to ensure maximal comparability across statistical classes. Figure 3B illustrates the iterative evaluation of Bayesian inference (*A*) and optimization (*I*_max<_) as the population dynamics unfolds in rescaled time *τ* (measured in the units of selection). Accuracy *A* may decrease significantly over time as the population, composed of particles with only local information, tracks an ideal posterior distribution with optimal global reallocation of probabilities. We investigate this phenomenon in detail below.

## Results

### Approximating the Bayesian posterior

To understand when and why populations estimate the true Bayesian posterior well, we first map evolutionary parameters, mutation rate per bit *m* and selection strength per generation *s*, onto the population summary statistics, mean information gain *μ*_*I*_ and standard deviation of information gain σ_*I*_ (see Figure 4A). Then, on Figure 4B, we show the relationship between summary statistics *μ*_*I*_ and σ_*I*_, and accuracy *A* of the population-based estimate of the true Bayesian posterior. In all panels of Figure 4, population size is set to *n* = 100. Although the parameter space of *μ*_*I*_ and σ_*I*_ is narrowly explored even when evolutionary parameters *m* and *s* span multiple decades, high accuracy (*A* ≈ 0.2) Bayesian inference is relatively easy to achieve on short timescales (*τ* = 1). However, at long timescales (*τ* = 10), Bayesian inference is not highly accurate because the population is unable to explore *most* high-fitness (high information gain) genotypes because they are scattered across the landscape. The timescale of environmental change is therefore a strong determinant of the computational capacities of Bayesian-Darwinian populations even when parameters such as mutation rate *m* and selection strength *s* cover a wide range from very low correlation between generations to high-fidelity replication, and from weak to strong selection.

**Figure 4.**
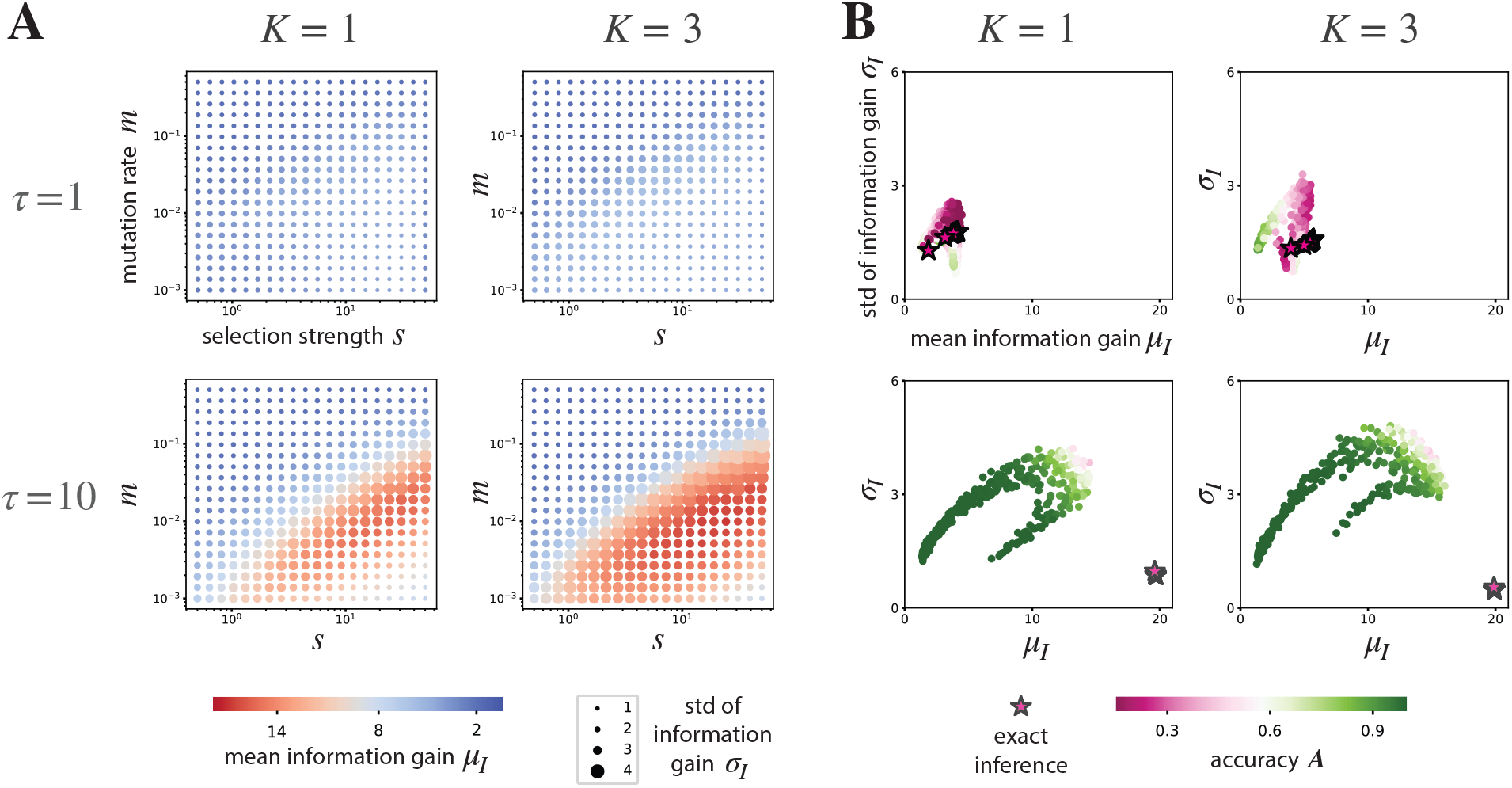
Effective optimization over long timescales vs effective estimation of the Bayesian posterior over short timescales. A) The population mean information gain *μ*_*I*_ (color) and its standard deviation σ_*I*_ (dot size) on the likelihood landscape, *P*_likelihood_ ∝ exp(*τ*αΦ_likelihood_), as a function of mutation rate *m* and selection strength *s*. B) *μ*_*I*_ and σ_*I*_ is further mapped to the accuracy *A* of the estimation of Bayesian posterior. High accuracy Bayesian inference can be realized by various *μ*_*I*_ and σ_*I*_ population measures, reflecting a near-optimal arbitration between remembering the past and fitting the present, encoded by the prior and the likelihood, respectively. In comparison, stars indicate the exact posterior *P*_posterior_ under various selection strength parameters *s*; they, by definition, correspond to perfect accuracy, *A* = 0. Under wide parameter regimes of dynamical parameters *m* and *s*, high accuracy Bayesian inference can be achieved over short timescales (*τ* = 1) and high mean fitness can be reached over long timescales (*τ* = 10), but not vice versa. On both panels, population size is *n* = 100, and samples are constructed from *n*_action_ = 100 runs.

### Adaptive hyperparameter trajectories

We further investigate the direct relationship between the hyperparameters mutation rate *m*, selection strength *s*, and population size *n*, on the accuracy *A* of Bayesian inference by the population. Since *m,s*,and *n* are all assumed to be evolvable at the level of collectives, adaptive paths in their 3-dimensional space could correspond to possible evolutionary scenarios. We focus on two main questions. *(i)* What is the smallest population size and the highest mutation rate (lowest information transmission) where a gradient in hyperparameter space is already present, driving the system towards lower mutation rate and larger population size? (*ii)* Can a population, with hyperparameters *m, s*, and *n*, evolved for Bayesian inference, be used as an efficient optimizer in combinatorial spaces? In other words, can performance in Bayesian inference, as a computational trait of a collective, serve as a preadaptation to combinatorial optimization?

As Figure 5 shows, a hyperparameter gradient is already present at *n* = 2,*m* ≈ 0.1 if selection is imposed by Bayesian inference at the level of collectives. This contrasts with the neutral landscape observed at *n* = 2 under selection by combinatorial optimization. Following a gradient on the Bayesian objective bridges serial (*n* = 2, one comparison per generation) to population-based (*n* ≈ 10) computation, while also lowering the mutation rate and increasing the number of generations *T*, to approximately *n,m,T* ≈ 10,0.03,10. After this Bayesian regime, both the Bayesian and the optimization objective leads to a further increase in population size and decrease in mutation rate, perfecting the system in both, achieving a functional flexibility where the function is determined only by the effective timescale of likelihood integration *τ*. This implies that efficient population-based combinatorial optimizers could evolve along an adaptive trajectory *only* on a Bayesian objective, starting from serial, high-noise, low iteration number computation, *n,m,T* ≈ 2,0.1,2, to a population-based dynamics with low mutation rate and many generations, *n,m,T* ≈ 100,0.01,20, which could be called a Darwinian regime. Taken together, this adaptive path leads to a “free” Darwinian combinatorial optimizer when the same dynamics is employed on longer timescales (*τ* = 10 vs 1), a functional exaptation of Bayesian inference.

**Figure 5.**
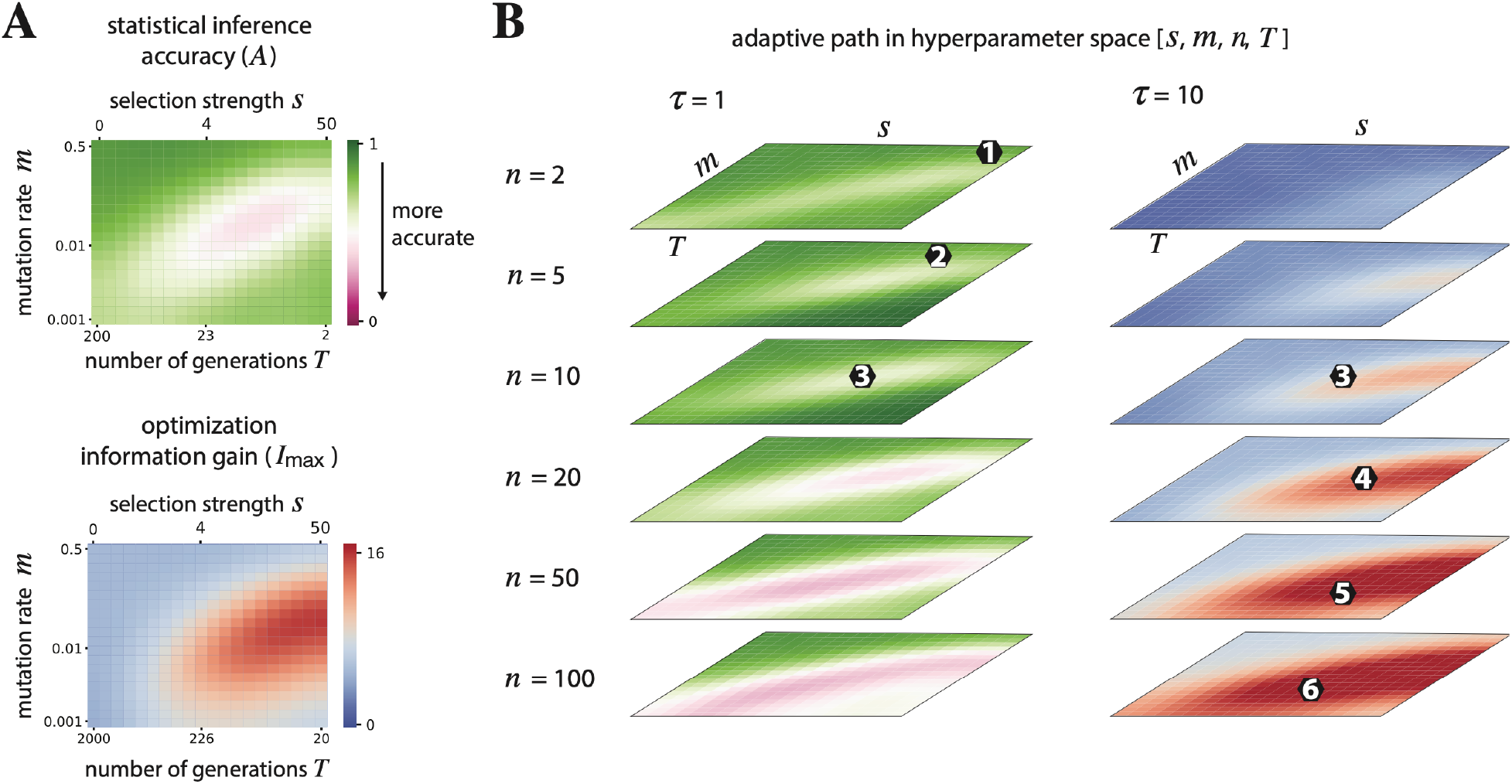
Darwinian coadaptation of Bayesian inference. Adaptive trajectory in hyperparameter space (*s,m,n,T*)connects a serial (*n* = 2, one comparison) sampling-based approximation of the Bayesian posterior, with low inherited information between subsequent samples (high *m*), to Darwinian dynamics of a large population (large *n*, low *m*) that performs well on a combinatorial optimization task (steps *1-6*). The Bayesian objective (left column) selects for more inherited information (lower *m*) before the combinatorial optimization objective (right column) does, bridging the critical gap from serial (*n* = 2) to population-based (*n* ≈ 10) computations. Further hyperparameter evolution on either objective leads to further decrease in mutation rate and increase in population size, a regime where the same dynamics can be flexibly employed to compute Bayesian inference or combinatorial optimization, set only by the timescale of the process *τ*. The results shown here correspond to a rugged landscape with *N* = 20 and *K* = 3.

## Discussion

Consider a humble prokaryote, say *Escherichia coli*. It feeds mainly on sugars, with glucose as its preferred carbon source. Changes in the concentration of food trigger movement orientation, so that *E. coli* swims in the direction it perceives to be most favorable, and otherwise changes direction. This glucose-seeking behavior, as a way of improving fitness, has been built up through evolution by fine-tuning genetic networks to regulate gene expression and update the information these organisms receive from a noisy environment. Of course, *E. coli* is no intellectual giant, but as Simpson (47) (p. 17) put it: “All the essential problems of living organisms are already solved in the one-celled … *creatures* and these are only elaborated in man or the other multicellular animals.” (We have replaced “protozoan” in the original by *creatures*.)

The former is just one example of how genetic evolution enables cells to perform sequential Bayesian inference (48–53). Cells live in a stochastic environment; for example, they must decide whether to turn on or off the biosynthesis of an enzyme that converts a substrate into a product. Typically, cells import the molecule of interest, measure its intracellular concentration to infer its extracellular concentration, and decide accordingly. These processes require autocatalytic components, sigmoidal response curves (48,50,53) and memory, implemented for example by specific receptors for particular signals in the cell membrane. This capacity has been co-opted by innate immunity, in which natural killer cells and macrophages possess the type of cellular reactivity and memory discussed above and use it to respond to invading challenges. A Bayesian description of this capacity (54) seems to explain several interesting dynamical facts, including a trade-off between precision and adaptation. Remarkably, this system seems to learn what is self, and attacks everything else that does not look like itself. Note that adaptation occurs at the cellular level, and the population level is just a direct outcome of the former. The adaptive immune system of vertebrates can do more: it uses a generative grammar and clonal selection to build up a vast antibody repertoire against invading antigens. This system learns about antigens directly through variation and selection, and can also be characterized as a Bayesian inference device (55). The Bayesian behavior of the population rests on the analogy between the Bayesian update rule and replicator dynamics. An explicitly Bayesian model of the functioning of memory within the adaptive immune system shows how an optimal balance between remembering old and collecting new pathogen evidence can minimize the risks of future infections (56).

Eukaryotes also evolved another sophisticated and radically different new way of sensing and responding to their environment: action potentials, a type of electrical signal that first evolved during eukaryogenesis (57). Later, specialized cells called neurons, initially organized into nerve networks that form the entire nervous system of some cnidarians (jellyfish), served to coordinate vital activities. After several hundred million years, intelligent creatures with large brains appeared. Action potentials are the electrical signals by which the brain receives, analyses and transmits information. Why has such a complex and energetically costly organ evolved? Despite the vast evolutionary distance between our humble *E. coli* and the arrival of animals with big brains, there is a common theme: both the food-seeking behavior of *E. coli* and a brain that helps animals respond to the demands of their environment can be understood as reward-based decision-making systems. Reward-based decision-making can be well represented as Bayesian probabilistic inference (58). Reward-driven motivation provides information that helps organisms increase their fitness. For example, implanting electrodes in the reward areas of rats promotes sexual behavior or any other naturally rewarding activity. Bumblebees have a greater response after conditioning with an increased sucrose concentration compared to an increased volume of sucrose solution for a similar total reward. This seems to be an adaptive behavior, as the nectar volume of a flower varies with foraging activity, whereas sucrose concentration is a reliable trait of the species (59). As a final example, magpies, which are birds in the Corvidae family and widely regarded as intelligent creatures, respond to visual cues that reliably indicate the presence of food and ignore less reliable spatial cues (60).

How the brain’s modulatory systems involved in reward, attention and motivation interact with the sensory and motor systems is one of the most exciting questions in neuroscience, and one that is fundamental to our understanding of learning and memory (61). One of the reasons we are beginning to understand how the brain’s reward system works is that it is similar to the method of reinforcement learning developed in AI, for which we have a very solid theory (62). For example, reward prediction error - the reduction in neuron firing when a predicted reward does not occur - was modelled using equations based on reinforcement learning theory (63).

Reinforcement learning uses only a positive reward or negative punishment to indicate whether the task is being performed correctly. It has found widespread and successful application in both cognitive science and AI, such as DeepMind’s AlphaGo.

From an evolutionary perspective, our model can be thought of as an approach where adaptive agents, and thus their reward functions, are evaluated according to a Bayesian optimisation framework that not only increases expected fitness but can also increase complexity. Statistical inference is a ubiquitous functional adaptation of animal minds that compete to predict changing environments (64,65). Better extraction of relevant statistical features from the past and optimal integration with current noisy sensory input allows for faster and better action selection. The first link to evolutionary dynamics is the realization that the Bayesian inference rule is a special case of the discrete-time replicator equation. This in itself suggests that an appropriate algorithm with an important evolutionary component can be expected to perform Bayesian inference as well as other tasks beyond classical Bayesian capabilities. The latter is justified by the greater scope of the replicator equation (for example, it can implement cooperation and competition between hypotheses by design). Conversely, existing algorithms that approximate Bayesian solutions have an interpretation in terms of evolutionary dynamics (3,4,7). Furthermore, our framework applies not only from a phylogenetic perspective, but also during ontogeny, as in the case of the immune system discussed above. Another well-established example is the relationship between Bayesian cognitive models and Darwinian neurodynamics (28,29,31), which occurs within a lifetime.

Darwinian neurodynamics posits that it can pay to generate and propagate candidate solutions through a *bona fide* evolutionary process in the brain (28), exploiting known component mechanisms, including massive parallelism (e.g. in the form of mini-columns), competition for cortical space, and reward allocation (31). The value of this hypothesis lies in its ability to suggest convincing and robust information transfer (i.e. some form of copying) between the parallel units, which is a hot topic of current research. Darwinian neurodynamics describes an effective parallel, stochastic search with redistribution of resources (by information transfer between the parallel units, based on the performance of their contents) (28). Although the application of the NK model in the present paper does not refer to a specific task, it is useful because, at an abstract level, it can correspond to the learning landscape on which the stochastic search in the language of thought is carried out (66–68). The proposed mechanism in this paper suggests a possible adaptive evolutionary route to the algorithmic emergence Darwinian neurodynamics. This mechanism complements previous work on *(i)* alternative models of neural information copying (29,31,69–71), *(ii)* experimental findings suggesting that *in vitro* information transfer between neural ensembles can emerge without explicit training (72,73), and *(iii)* proposed cognitive markers of a Darwinian search in hypothesis space (30).

One might object that classical reinforcement learning (RL) could do the job that Darwinian neurodynamics is supposed to do. It has been repeatedly pointed out that Skinnerian operant conditioning is loosely analogous to a selection process (33). Do we need more? RL struggles with the complexity posed by problems such as diverse exploration, conflicting reward goals, sparse rewards, reward shaping, hyperparameter tuning, and policy search in large combinatorial spaces (74). Various evolutionary approaches (including evolution strategies, genetic algorithms, and genetic programming) have successfully been applied to augment basic RL methods (74).

This strengthens the view that the phylogenetic progression has led from Bayesian to Bayesian-Darwinian brains in evolution, reaching significant levels of performance in cephalopods, birds and mammals. In the hominin lineage this mechanism is hypothesized to have evolved open-ended exploration-exploitation capacity.

The evolution from a Bayesian to a Bayesian-Darwinian system may play a crucial role in the spontaneous or designed emergence of Darwinian evolution across very different physical implementations. Complex small-molecule autocatalytic systems exhibit remarkable adaptations to environmental conditions (75), as predicted by theory (76,77). A paradigmatic example is the formose reaction, which is based on the autocatalytic formation of sugar molecules from a formaldehyde source. The network contains a number of autocatalytic sub-cycles that cross-react in various ways, but does not exhibit hereditary variation over cycles. Nevertheless, adaptive tracking of the environment is remarkable; shifts in environmental variables are matched by internal rearrangements of concentrations and fluxes, corresponding to altered dominance of the embedded sub-cycles. This capacity points to an intriguing fundamental question: how could complex autocatalytic systems have evolved first limited, and then unlimited heredity (23,24). Accurate replication emerged during the origin of life process on Earth. Modelling of this evolutionary process in the form of the Bayesian to Darwinian agents is a subject for further study. In particular, modelling of primordial, reflexively autocatalytic peptide network (43) evolution (24) would allow for the use of the NK landscape to develop the investigations on this present paper further.

We also briefly note that the relationship between evolutionary dynamics and various Bayesian approaches is an area of active research. The isomorphism between Bayesian updating and the discrete-time replicator equation (3) is now a widely known starting point. Frank (78) analyzed some fundamental equations of evolutionary dynamics from the point of view of information gain and offered a Bayesian interpretation of selection. These approaches seem obvious with hindsight, since every allele is a hypothesis about how to fare in the given environment, and it is not surprising that bad ideas are removed by natural selection. Vanchurin et al. (79) portray evolution as multilevel learning; the derivation rests on renormalization and separation of timescales. Importantly, they start with learning and derive evolution from that, rather than the other way around. A related approach (80) unites intra- and intergenerational change through a variational Bayesian approach and arrives at a unified equation resting on the separation of timescales, renormalizability, variational free energy and the principle of least action, under the assumption that a non-equilibrium steady state distribution exists. The variational free energy has been applied to the problem of natural language origins (81) in a population of agents who profit from sharing beliefs, given a shared world model. We note that the latter is analogous to information exchange by sexual recombination allowed by a shared genetic code in the interacting agents. We emphasize that the present paper takes an explicitly mechanistic approach, following the dynamics of a population by numerical simulation. A more abstract formal treatment seems feasible in the future, considering the work in the previous paragraph.

## Code availability

Code is available at https://github.com/csillagm/BayesToDarwin.

## Author contributions

Conceptualization: HG, ES, DC. Model: MC, HG, DC. Simulations: MC. Figures: MC, HG. Final manuscript: MC, HG, ES, MS, DC.

## Acknowledgements

We thank the two anonymous reviewers for their helpful comments on the manuscript. We acknowledge financial support from the Hungarian National Research, Development, and Innovation Office (NKFIH, grant KKP129848, MC, ES, DC), and the Templeton World Charity Foundation (grant TWCF0268, ES, DC), as well as the MiniLife ERC Synergy Grant (101118938, ES, DC). MS is supported by grants PID2021-127107NB-I00 from the Ministerio de Ciencia e Innovación (Spain), 2021 SGR 00526 from Generalitat de Catalunya 2021 SGR 00526, and the Distinguished Guest Scientists Fellowship Programme of the Hungarian Academy of Sciences.

